# From bones to sediments: ancient human DNA from open-air archaeological sites

**DOI:** 10.1101/2025.03.19.643861

**Authors:** Rikai Sawafuji, Ryohei Sawaura, Masaki Yokoo, Toshiaki Kumaki, Nils August Thomasen, Takumi Tsutaya, Mikkel Winther Pedersen

## Abstract

Ancient human bones degrade and sometimes disappear, especially in regions with acidic soils, making it difficult to obtain genetic information from past populations. To explore alternative sources, we analyzed ancient DNA from sediment samples collected from a burial site and a settlement site in Japan, both dating to around 1000 years ago. We found that ancient human mitochondrial DNA was obtained from sediments surrounding bones, particularly close to rib bones, while human DNA was rarely detected in the settlement site sediments. Furthermore, the mitochondrial haplogroups identified in the sediments were identical to those from human bones, confirming the reliability of this approach. Our findings suggest that genetic information about past human populations can be directly obtained from archaeological sediments in open-air sites. This method also provides a non-destructive alternative to bones and teeth, expanding possibilities for ancient DNA research in regions where skeletal remains are poorly preserved.

## Introduction

The analysis of ancient human DNA has revolutionized our understanding of human migration and dispersal, reshaping our historical narratives with unprecedented detail (Allentoft et al., 2015; Haak et al., 2015; Moreno-Mayar et al., 2018). However, the field remains affected by substantial biases and skewness, particularly in geographical and temporal representation. This is in part due to the disproportionate allocation of research funding and archaeological activity by the Global North. Furthermore, climatic and environmental factors contribute to this imbalance, as fossil preservation is more favorable in cooler, stable environments, thereby limiting the recovery of genetic material from regions with less suitable preservation conditions (Allentoft et al., 2012; Emmons et al., 2020; Eriksen et al., 2020). In such regions, bones may undergo complete degradation, resulting in the absence of recoverable remains at sites. Even when skeletal remains are discovered, endogeneous DNA is often highly degraded. Consequently, ancient human DNA sampling is disproportionately concentrated in regions with more favorable preservation conditions and where research efforts are most intensive. This concentration results in a skewed understanding of human migratory and dispersal patterns, overlooking key areas where hominins would have interacted and evolved (Gokcumen & Frachetti, 2020).

In light of these challenges, the analysis of ancient human DNA from sediments has emerged as a promising alternative. For instance, ancient hominin mitochondrial DNA (mtDNA) from Denisovans and Neanderthals has been successfully recovered from sediments in Denisova Cave (Slon et al., 2017; Zavala et al., 2021) and Baishiya Karst Cave on the Tibetan Plateau (Zhang et al., 2020). Furthermore, nuclear genomes have also been extracted and analyzed from cave sediments (Gelabert et al., 2021; Vernot et al., 2021). These new approaches expand the scope of hominin ancient DNA research beyond skeletal remains, offering a novel method to investigate the history of human presence and migration (Sawafuji et al., 2024).

Although ancient human DNA in sediments holds great potential, significant challenges still limit its utility. A primary issue is identifying which specific locations and contexts within archaeological sites are most likely to yield ancient human DNA. From the few published studies, the success rate of recovering human DNA from sediments seems highly variable and influenced by factors such as stratigraphic layers, mineral composition, and sites-specific conditions (de-Dios et al., 2024; Massilani et al., 2022; Zavala et al., 2021). Nevertheless, more studies comparing the extent of human DNA recovery across different archaeological contexts have yet to be conducted, underscoring the need to validate and improve the success rates of sediment-based DNA analysis across diverse sites, climates and timescales.

Even more pronounced is the lack of studies on human DNA in sediments from open-air archaeological sites, which remain less explored compared to cave or sheltered contexts. Human DNA to date has been successfully obtained from calcite stones covering skeletal remains in a cave (Sarhan et al., 2021), as well as from sediments in caves (Gelabert et al., 2021; Massilani et al., 2022; Slon et al., 2017, 2022; Vernot et al., 2021; Zavala et al., 2021; Zhang et al., 2020) and beneath a rock shelter (Zampirolo et al., 2024). In contrast, open-air sites, which have the potential to provide a broader range of ancient human DNA, remain significantly underexplored. Given that the number of open-air sites with evidence of human activity far exceeds that of cave sites with good preservation, future studies exploring the potential of sites with poor or no bone preservation are crucial for expanding the diversity of ancient human genetic studies and our history.

Japan has a diverse climate, spanning multiple climatic regions, but high humidity and acidic soils across much of the country contribute to poor bone preservation, making ancient skeletal remains scarce (Gakuhari et al., 2020). This phenomenon is not unique to Japan but is also common in other regions where similar environmental conditions prevail, such as Southeast Asia (McColl et al., 2018). Due to these poor preservation conditions, a comprehensive understanding of prehistoric human presence and dynamics remains challenging. Despite this, Japan’s wide variety of archaeological sites provides a valuable case for testing sediment-based DNA recovery techniques. Extracting DNA from sediments could offer a crucial alternative for reconstructing ancient human history and serve as a model for other regions with similar preservation challenges.

In this study, we evaluate the potential for recovering human DNA from sediments at two open-air archaeological sites in Japan, a burial site and a dwelling site, using shotgun sequencing and mitochondrial DNA target capture enrichment.

## Results

Five sediment samples were collected at a burial site, Katsuren castle, located in Okinawa, which dates to the mid-12th to early 13th century during the Gusuku period (Figure 1). Here, two burials with skeletal remains were found beneath the castle’s wall. One sediment sample from inside the skull was collected from Pit 138. From Pit 178, four sediment samples were obtained: one from inside the skull, one from sediment around the hip bones, one from sediment around the rib bones, and one from an area outside the pit (Figure 1, Table S1). Additionally, one rib bone was collected from each individual to compare with the results obtained from the sediments. The second site is a dwelling site known as Oshima 2, which dates to the 11th to 12th century during the Satsumon period and is located in Hokkaido (Figure 1). We collected eleven sediment samples from different contexts within the site (Figure S1, Table S2). DNA was extracted from all samples and sequenced using Illumina NovaSeq and iSeq platforms (see Methods). We performed a combination of shotgun sequencing (between 12447839 and 85971229 number of reads) and targeted capture enrichment for human mtDNA (between 70064 and 2174014 number of reads) to investigate the feasibility of determining human mtDNA haplogroups from sediment samples in open-air archaeological sites.

**Figure 1.**
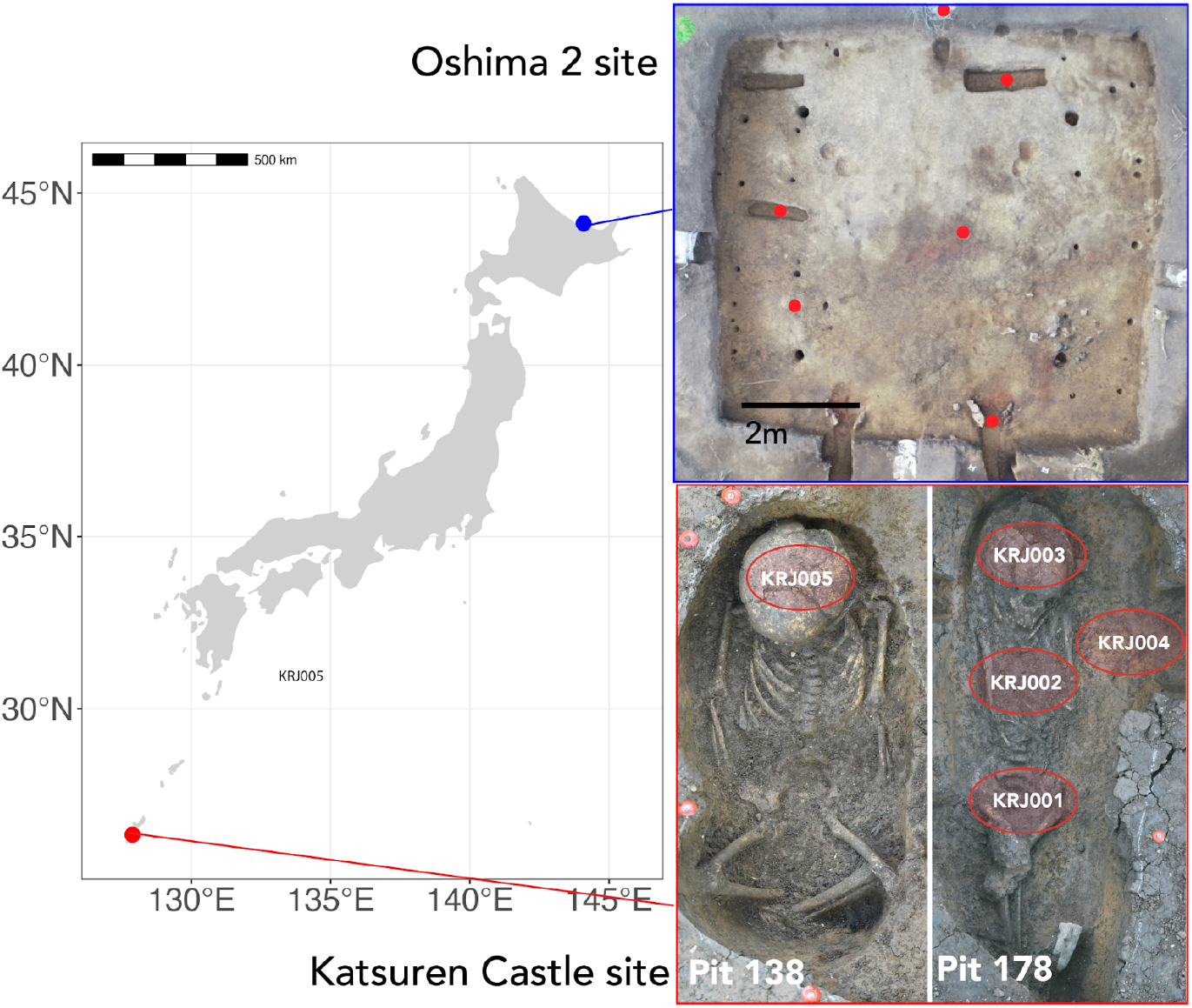
Locations of two sites. Red dots and circles indicate the locations of collected sediment samples.

Haplogroups were successfully identified from sediment samples at the burial site after mitochondrial DNA target capture (Figure 2, Table S3). Ancient DNA authenticity was confirmed by assessing ancient DNA damage patterns such as thymine misincorporation from cytosine deamination (Figure S2) and by analyzing the distribution of read length (Figure 2a) and damaged reads (PMD score > 2) across the reference genome (Figure 2b). Haplogroups identified in the sediment samples matched those from the bone remains from the same pits: M7a1b1 from Pit 138 and D4a1 from Pit 178. Specifically, one sediment sample from each pit yielded mtDNA haplogroups. In contrast, sediment samples from the dwelling site did not yield sufficient human mtDNA for haplogroup determination or damage pattern analysis, even after capture enrichment (Figure S3).

**Figure 2.**
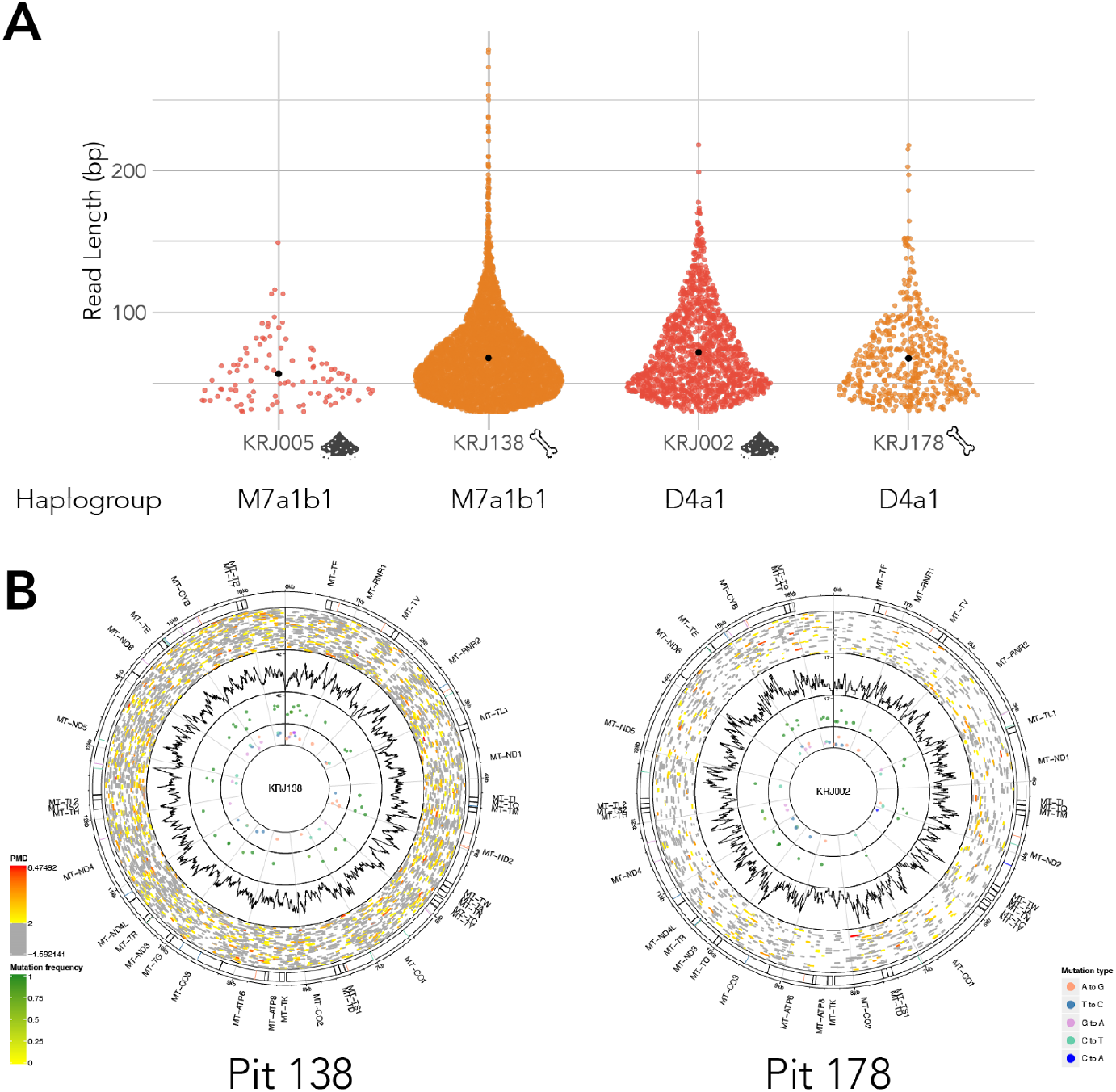
MtDNA read length and its haplogroup. A) Read lengths of mitochondrial DNA and their haplogroups. B) Distribution of the mapped reads.

We also analyzed the shotgun sequencing data from sediments and bones. Samples with determinable haplogroups displayed clear damage signatures when mapping to the human nuclear and mitochondrial genome using shotgun datasets, while samples without identifiable haplogroups did not (Figure 3, Figure S3). However, shotgun sequencing alone yielded insufficient reads for haplogroup determination, sex determination, or PCA. Targeted capture significantly improved mtDNA recovery, increasing coverage depth 30-to 200-fold in samples with identifiable haplogroups, with the greatest increase observed in sample KRJ002 (Figure S4).

**Figure 3.**
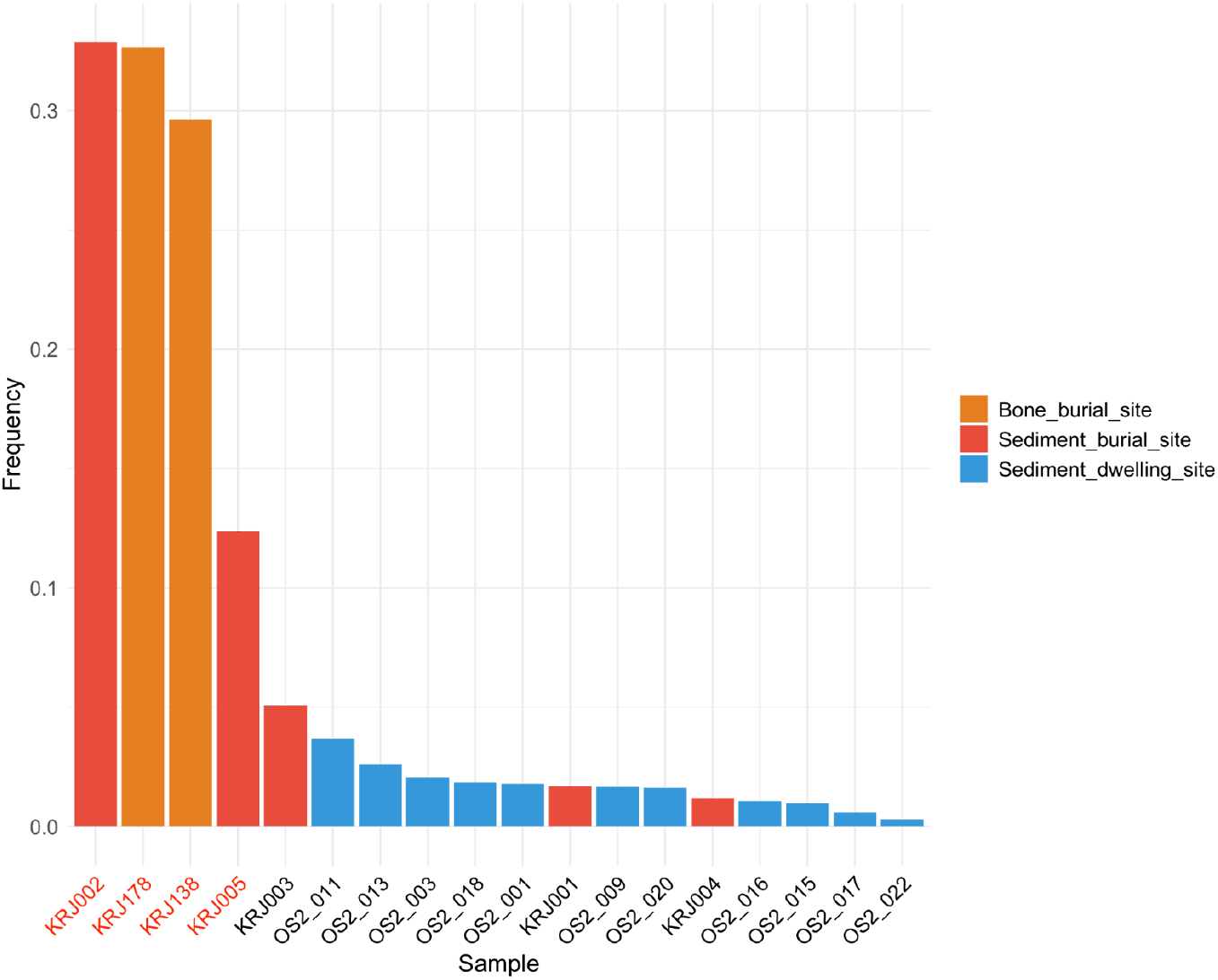
Damage degree of shotgun reads mapped to human genome. The red text color indicates samples that the haplogroup was determined from.

To assess DNA degradation, we analyzed the bacterial DNA fraction from the shotgun datasets of sediments and bones. Most samples, except for KRJ001 and KRJ005, showed highly fragmented bacterial DNA, with average fragment lengths below 60 bp (Figure S5). KRJ001 and KRJ005 from the burial site contained relatively longer DNA fragments (>100 bp), suggesting possible intrusion by modern bacteria. While all samples showed bacterial taxa with damage patterns (Figure S6), these were more prevalent at the burial site, indicating poorer DNA preservation compared to the dwelling site. Principal Coordinate Analysis (PCoA) of the bacterial microbiomes indicated a tendency toward site-specific similarities. (Figure 4). Notably, at the burial site, bacterial communities from the bones and the sediment from inside the skull (KRJ005) were more similar to each other than to the bacterial communities in other sediments.

**Figure 4.**
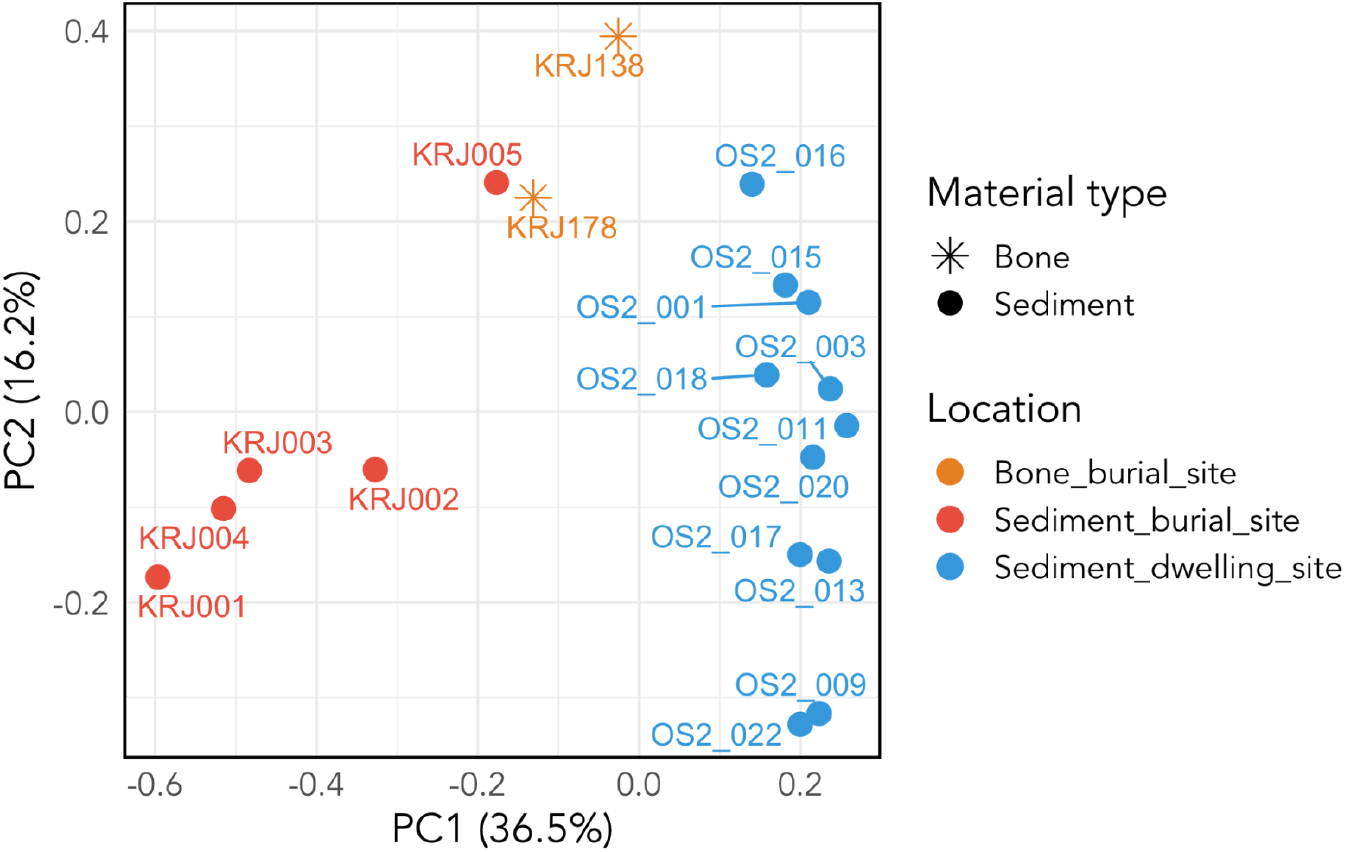
PCoA analysis using bacteria dataset.

## Discussion

This study presents two significant findings: (1) ancient human mtDNA can be successfully recovered from open-air archaeological sites, and (2) the abundance of human mtDNA varies considerably based on the site’s location and context. This highlights that human activity at an archaeological site does not necessarily ensure the recovery of ancient DNA or ancient human DNA from sediments.

Our results mark an early foundational step in exploring human DNA recovery from open-air archaeological contexts. By recovering human mtDNA from both bones and sediments and identifying matching haplogroups with characteristic degradation patterns, we provide a robust framework for future sediment-based human DNA analysis. These findings highlight the potential to extend human DNA analysis to a broader range of open-air sites, providing a foundation for broader applications.

### Contextual variation in DNA recovery

The recovery of human DNA varied depending on the archaeological context. Burial sites are likely the most promising, as ancient human mtDNA was recovered from sediments surrounding the bones, particularly near the rib bones. This aligns with a previous finding from a cave site (Sarhan et al., 2021) and suggests that the presence of decomposing remains contributes significantly to human DNA recovery in sediments.

In contrast, human DNA was rarely detected in sediments from the dwelling site. This is likely because the primary candidates of human DNA resources in this context are hair, skin and sweat, which are typically present in low quantities and may not yield enough DNA for ancient human DNA analysis. Additionally, limited direct contact between inhabitants and the ground due to footwear use may have further reduced the deposition of human DNA. Further studies are required to confirm the absence of human DNA in sediments at dwelling sites and explore other contexts such as latrines and middens, which have yielded DNA in previous studies (Sabin et al., 2020; Søe et al., 2018). Such contexts may offer valuable DNA sources, especially if methods like microsampling or other improvements can isolate DNA from individual contributors.

### Insights from Microbial Analysis

Microbial community analysis provided additional insight into taphonomic processes. PCoA revealed a tendency toward differing bacterial compositions at the burial site compared to other sediment samples (Figure 4). Notably, the bacterial community in sediments inside the skull (KRJ005) closely resembled that of the bone, suggesting that the sediment DNA in this context reflects decomposition from human remains. This finding also highlights the dominant role of human tissues as a DNA source and underscores the need for further taphonomic studies to refine interpretations of DNA preservation and degradation.

### Implications for Screening and Ethical Considerations

This study highlights the potential of sedimentary ancient DNA (sedaDNA) as a valuable tool for screening ancient human DNA. Advances in DNA recovery from non-destructive materials like chewing gum (Jensen et al., 2019), dental calculus (Ozga et al., 2016; Ziesemer et al., 2019), and coprolites (Hagan et al., 2020) demonstrate the feasibility of extracting DNA from sediments. On one hand, such sediment-based recovery methods enable rapid screening during excavations without damaging human remains, supporting heritage preservation, and addressing some ethical concerns related to destructive analyses.

On the other hand, this approach introduces new ethical challenges, particularly when working with materials from contexts with sensitive historical, cultural, and social implications (Der Sarkissian et al., 2021). The lack of clear regulations or guidelines for handling ancient metagenomes compounds these concerns. Researchers must adopt responsible practices, engaging with stakeholders - including local communities and indigenous groups (Alpaslan-Roodenberg et al., 2021; Johnson et al., 2024) - to define project goals, manage sample storage, and communicate results transparently. Collaboration, transparency, and education on ethical practices must remain central to ancient metagenomic studies to ensure ethical and sustainable research.

### Limitations of the Study

This study examined sediments from only two archaeological sites dating to approximately 1000 years ago. The dwelling site, located in northern Japan (Hokkaido), generally exhibited better DNA preservation than the burial site, located in southern Japan (Okinawa), as indicated by bacterial DNA results (Figure S6). Despite this, no human DNA was detected from the dwelling site, suggesting a degree of robustness to our findings. Future studies should include a larger number of sites and various contexts to further refine sampling strategies. The coverage of the mitochondrial data obtained was insufficient for detailed kinship analysis or fine-scale haplogroup resolution. Additionally, our results are based solely on mtDNA; future work should aim to include nuclear DNA and sex determination.

## Materials and Methods

### Sample information

Sediments were collected from two archaeological sites in Japan (Figure 1). The first site, the Katsuren Castle site in Okinawa, is a burial site dating back to the Gusuku period, around 1000 years ago. Human skeletons were excavated from two separate pits, both located beneath the castle wall. Four sediment samples were obtained from Pit 178, and one from Pit 138 (Table S1). Additionally, rib bones from each individual buried in these pits were also sampled to verify whether the haplogroup of mtDNA from the sediment is identical to that from bone. The second site, the Oshima 2 site, Hokkaido, is a dwelling site dating back to the 11th to 12th century in the Satsumon period. The date of this site is estimated from the chronology of pottery types and radiocarbon dating from another dwelling in the same site. We took 11 sediment samples from pits, an oven, floors and outside the dwelling (Figure S1, Table S2).

### Radiocarbon measurement

Radiocarbon measurement was carried out by Palynosurvey Co., Ltd. at Yamagata University on the two human skeletal individuals from Pit 138 and Pit 178 of the Katsuren Castle site. Collagen was extracted from a left rib and the left tibia for skeletons from Pit 138 and 178, respectively, based on the method described in Tsutaya et al. (2017). Carbon and nitrogen stable isotope ratios were measured to estimate the marine reservoir effect using an elemental analyzer-isotope ratio mass spectrometry (EA-IRMS). Radiocarbon concentration was measured using an accelerator mass spectrometry (AMS). The radiocarbon age was calibrated against atmospheric and marine calibration curves IntCal20 and Marine20 (Heaton et al., 2020; Reimer et al., 2020) using the software OxCal, version 4.4 (Bronk Ramsey, 1995). We assumed that the contribution of carbon derived from marine products was 20 ± 10% based on the results of the stable isotope analysis (Table S4). By considering the local marine reservoir effect (⊿R = -143 ± 33, recalculated with Marine20) around Okinawajima Island (Yoneda et al., 2007), the 95% confidence intervals of the calibrated age were 1157–1279 cal AD for both skeletons (Table S4).

Radiocarbon measurement was carried out by Paleo Labo Co., Ltd. on three charred woods from the Oshima 2 site (Table S5). Charred wood samples were treated with the acid-base-acid method described in Hajdas et al. (2017) with slight modifications. Briefly, samples were washed with acetone and sequentially treated with 1.2 M HCl, 1.0 M NaOH, and 1.2 M HCl. Radiocarbon concentration for the resultant graphite samples was measured using an accelerator mass spectrometry (AMS) at Paleo Labo. The radiocarbon age was calibrated against an atmospheric calibration curve IntCal20 (Reimer et al., 2020), using the software OxCal, version 4.4 (Bronk Ramsey, 1995). The 95% confidence intervals of the calibrated age were middle 11th to early 13th centuries AD for all three samples (Table S5). The radiocarbon measurement was applied to the charred wood samples without a final growth ring, so the measurement results may have been affected by the tree rings that formed in older years. Therefore, the true age of the tree’s death or the site occupation would be slightly more recent than the obtained radiocarbon ages.

### DNA extraction, library construction, capture enrichment and sequencing

Approximately 150 mg of either sediment or powdered bone were extracted for DNA using the method described by Rohland et al. (2018). All libraries were constructed using the NEBNext Ultra II DNA Library Prep Kit for Illumina (New England Biolabs) following the manufacturer’s protocol. Libraries were amplified with 12 PCR cycles using Pfu Turbo Cx. Human mtDNA capture was performed using the myBaits Mito (Daicel Arbor Biosciences) following the manufacturer’s protocol v5.01. Sequencing was performed on the Novaseq and iSeq platforms for both shotgun sequencing and target capture sequencing. The raw sequencing data have been deposited at the European Nucleotide Archive (ENA) and are publicly available as of the date of publication (ProjectXX).

The sequence datasets, both shotgun and target-capture sequencing, were mapped against the human genome. We used fastp (Chen et al., 2018) to trim the adapters and low-quality sequences, and to merge pair-end reads with 11bp overlap. Sequences after these steps were aligned to the human reference genome and human reference mtDNA genome (rCRS) using bwa (Li & Durbin, 2009). Duplicate reads were removed using picard’s MarkDuplicates (http://broadinstitute.github.io/picard/). Subsequently, reads assigned to human mtDNA were extracted using SAMtools (Danecek et al., 2021), and haplogroups were determined using haplocart (Rubin et al., 2023) and haplogrep (Schönherr et al., 2023). We used DamageProfiler (Neukamm et al., 2021) to check the damage patterns of ancient DNA as an authentication and contamMIX (Fu et al., 2013) to estimate the level of contamination.

Microbiome analysis was conducted using a workflow described by Fernandez-Guerra et al. (2023) for shotgun sequencing data from all samples. We performed Principal Coordinate Analysis (PCoA) to examine the relatedness of the microbial compositions across the samples.

## Acknowledgments

This study was supported in part by Grants-in-Aid for Scientific Research (KAKENHI: 20H05822, 20J00078, 21K00989, 22KJ1413) from Japan Society for the Promotion of Science. We thank Naomi Doi for supporting this research and Kei Aono and Ayuko Kubo for their comments on the manuscript.

