## Supplemental Information for "From bones to sediments: ancient human DNA from open-air archaeological sites"

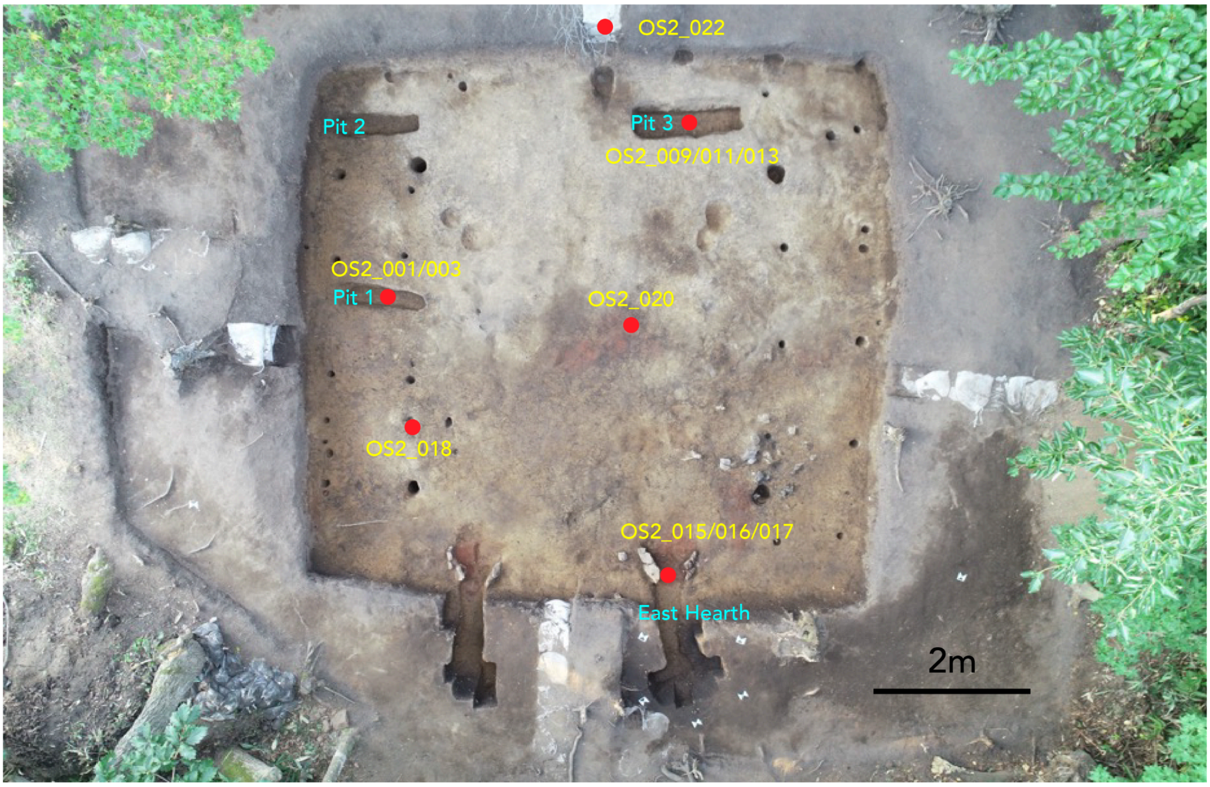


Figure S1. Sampling location in the Oshima 2 site.

Table S1. Sampling information from the Katsuren Castle site.

| Sample ID | Type | Pit | Sampling location |
|---|---|---|---|
| KRJ178 | bone | 178 | Rib bone |
| KRJ001 | sediment | 178 | Sediment around the hip bone |
| KRJ002 | sediment | 178 | Sediment around the rib bone |
| KRJ003 | sediment | 178 | Sediment inside the skull |
| KRJ004 | sediment | 178 | Sediment apart from the pit |
| KRJ138 | bone | 138 | Rib bone |
| KRJ005 | sediment | 138 | Sediment inside the skull |

Table S2. Sampling information from the Oshima 2 site.

| Sample ID | Type | Sampling location |
|---|---|---|
| OS2_001 | sediment | Sediment from the middle layer of soil deposition in the Pit 1 |
| OS2_003 | sediment | Sediment from the bottom layer of soil deposition in the Pit 1 |
| OS2_009 | sediment | Sediment from the upper layer of soil deposition in the Pit 3 |
| OS2_011 | sediment | Sediment from the bottom layer of soil deposition in the Pit 3 |
| OS2_013 | sediment | Sediment from the middle layer of soil deposition in the Pit 3 |
| OS2_015 | sediment | Sediment from the bottom layer of soil deposition in the East Oven |
| OS2_016 | sediment | Burnt sediment from the East Oven |
| OS2_017 | sediment | Sediment from the middle layer of soil deposition in the East Oven |
| OS2_018 | sediment | Floor surface 1 |
| OS2_020 | sediment | Floor surface 2 |
| OS2_022 | sediment | Outside of the dwelling site |

Table S3. Haplogroup from sediment/bone samples.

| Pit | Type | Sample ID | DNA length | coverage (x) | haplogroup |
|---|---|---|---|---|---|
| 138 | Sediment | KRJ005 | 56.8 | 0.34 | M7a1b1 |
| 138 | Bone | KRJ138 | 67.9 | 18.66 | M7a1b1 |
| 178 | Sediment | KRJ002 | 71.8 | 6.09 | D4a1 |
| 178 | Bone | KRJ178 | 67.7 | 2.08 | D4a1 |

Table S4. Stable isotopic and radiocarbon results of the subject skeletons excavated from the Katsuren Castle site.

| Sample | %C | %N | $\delta^{13}C$ | $\delta^{15}N$ | C/N | Element | $^{14}C$ age | Calibrated $^{14}C$ age (95% CI) |
|---|---|---|---|---|---|---|---|---|
| Bone from Pit 178 | 44.6 | 14.63 | -16.43 | 10.13 | 3.56 | Left tibia | 900 ± 20 | 1157–1279 cal AD |
| Bone from Pit 138 | 47.01 | 16.74 | -17.77 | 10.61 | 3.28 | Left rib | 900 ± 20 | 1157–1279 cal AD |

Table S5. Radiocarbon results of the charred woods excavated from the Oshima 2 site.

| Sample ID | $^{14}C$ age | Calibrated $^{14}C$ age (95% CI) |
|---|---|---|
| PLD-53730 | 920 ± 15 | 1041–1107 cal AD (56.98%)<br>1115–1176 cal AD (36.76%)<br>1194–1200 cal AD (1.72%) |
| PLD-53731 | 895 ± 15 | 1049–1081 cal AD (24.20%)<br>1134–1137 cal AD ( 0.47%)<br>1152–1218 cal AD (70.79%) |
| PLD-53732 | 945 ± 20 | 1035–1054 cal AD (14.75%)<br>1060–1157 cal AD (80.70%) |

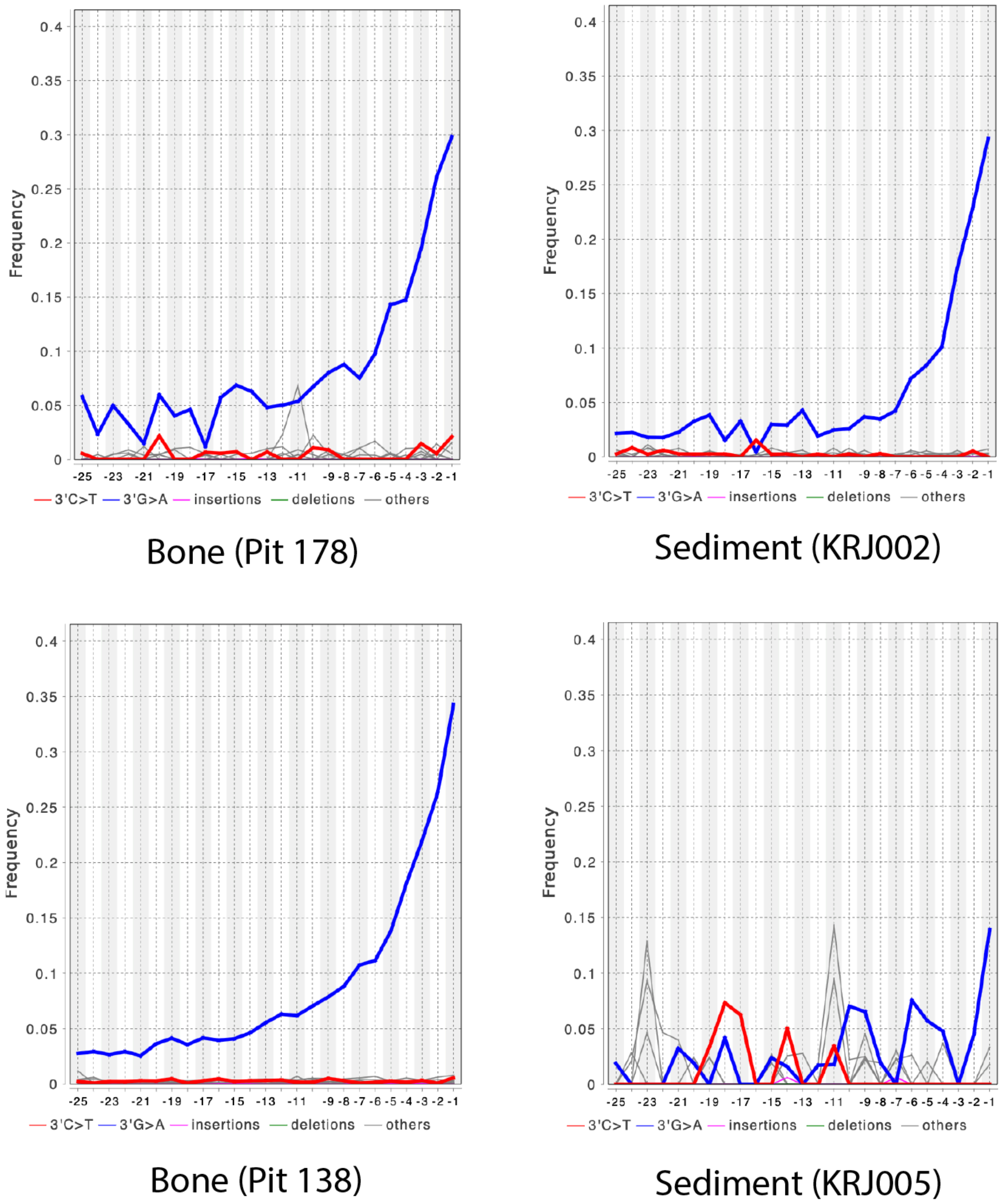


Figure S2. Damage patterns of human mtDNA from sediments and bones after enrichment.

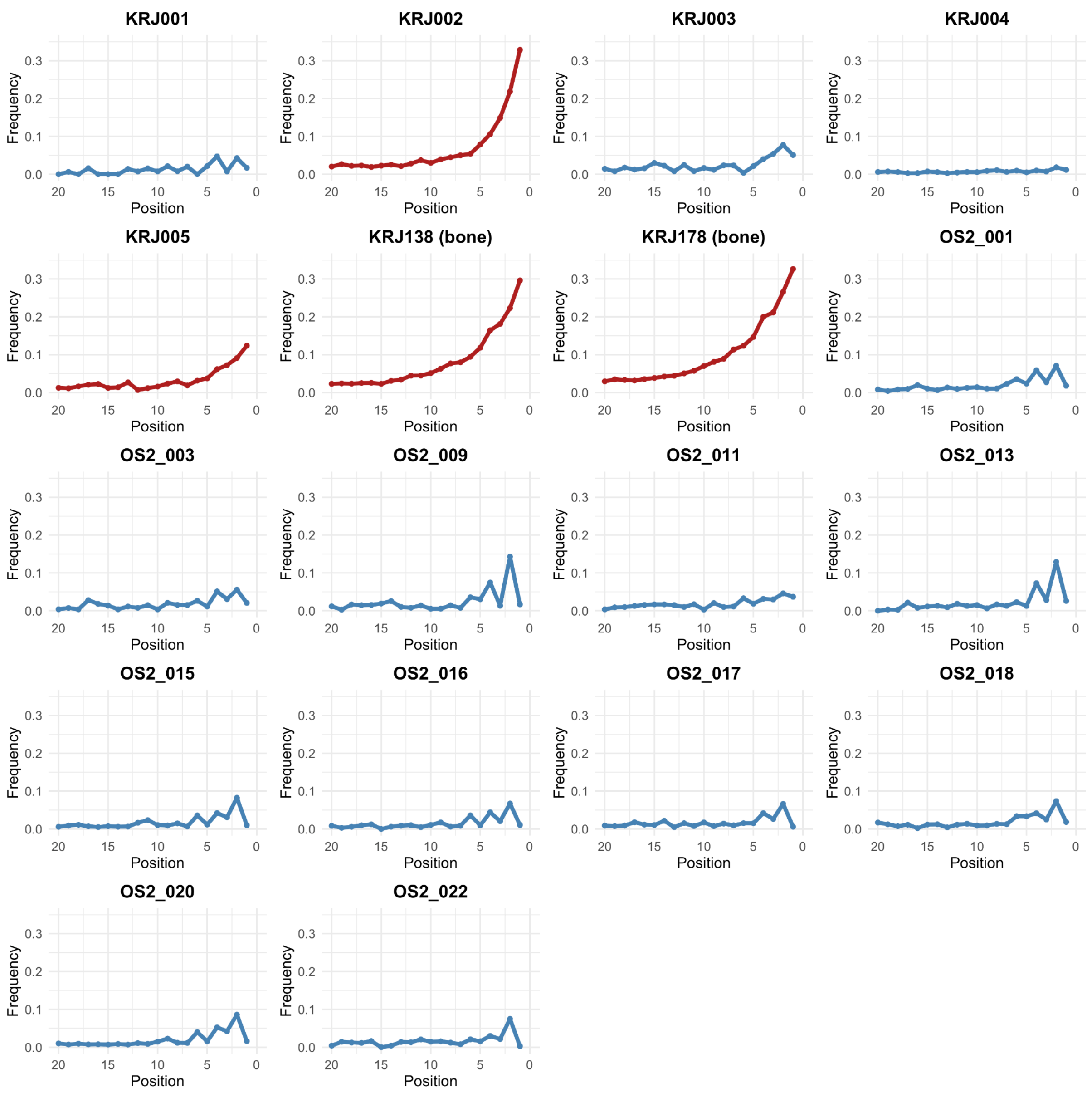


Figure S3. Frequencies of C to T transitions due to DNA Damage across the human nuclear and mitochondrial genome from both sediment and bone samples in the shotgun-sequenced datasets. Samples with the red line indicate successful enrichment of human mtDNA.

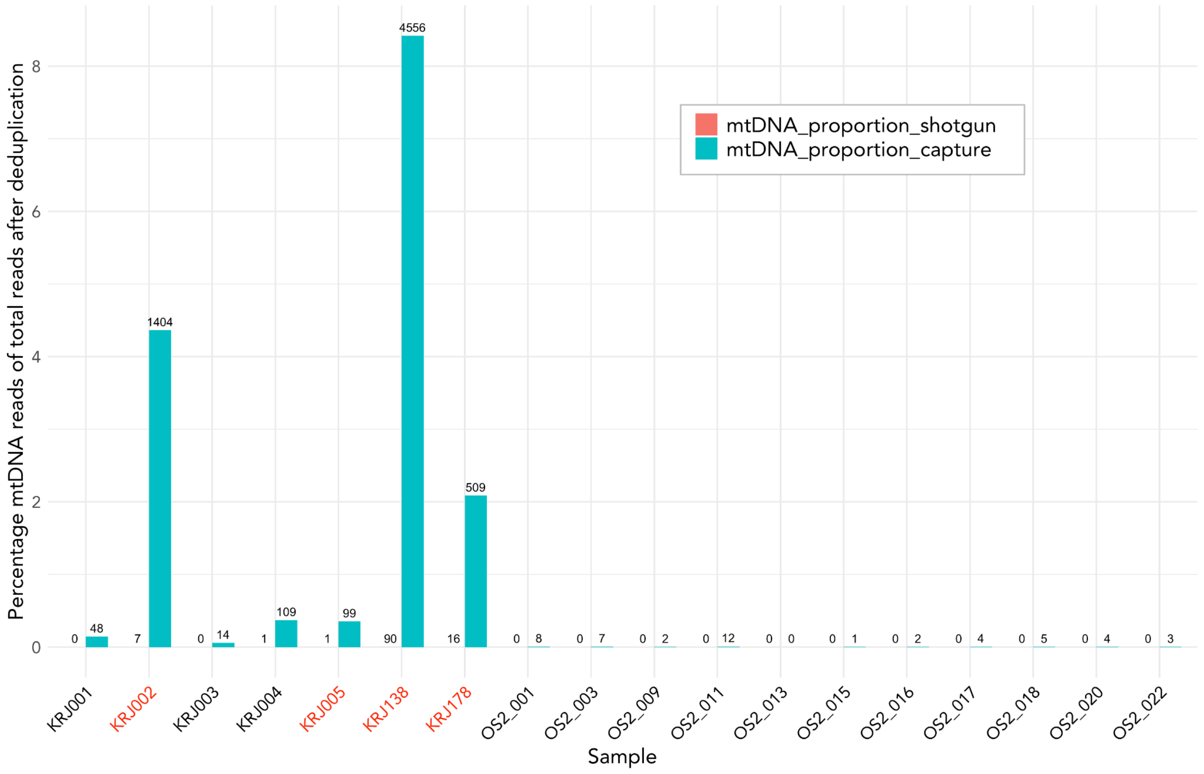


Figure S4. Capture efficiency of each sample. The red text color indicates samples that the haplogroup was determined from.

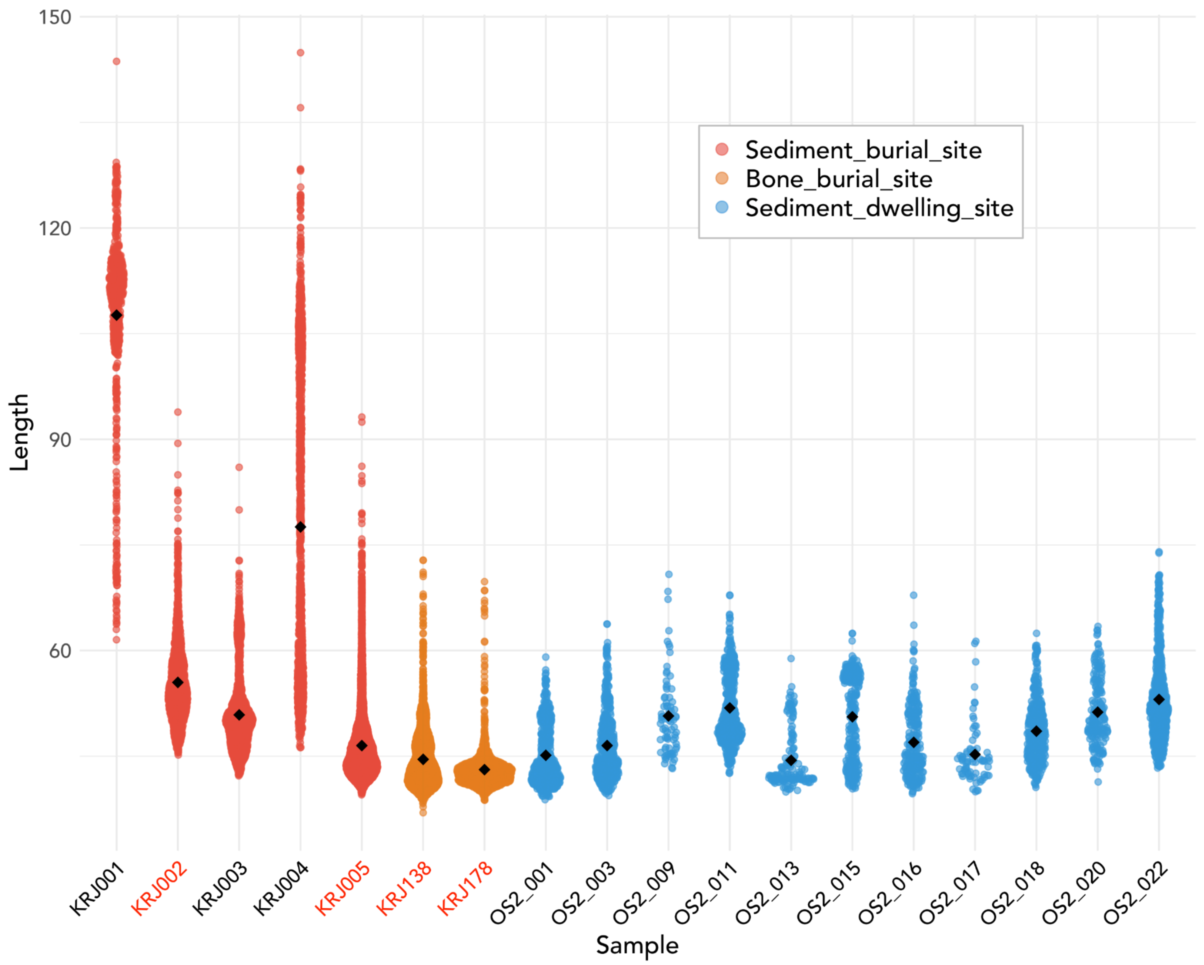


Figure S5. DNA length of each bacteria using shotgun datasets. Each dot represents the average DNA length of each taxon. The black dot represents the average DNA length of all taxon. The red text color indicates samples that the haplogroup was determined from.

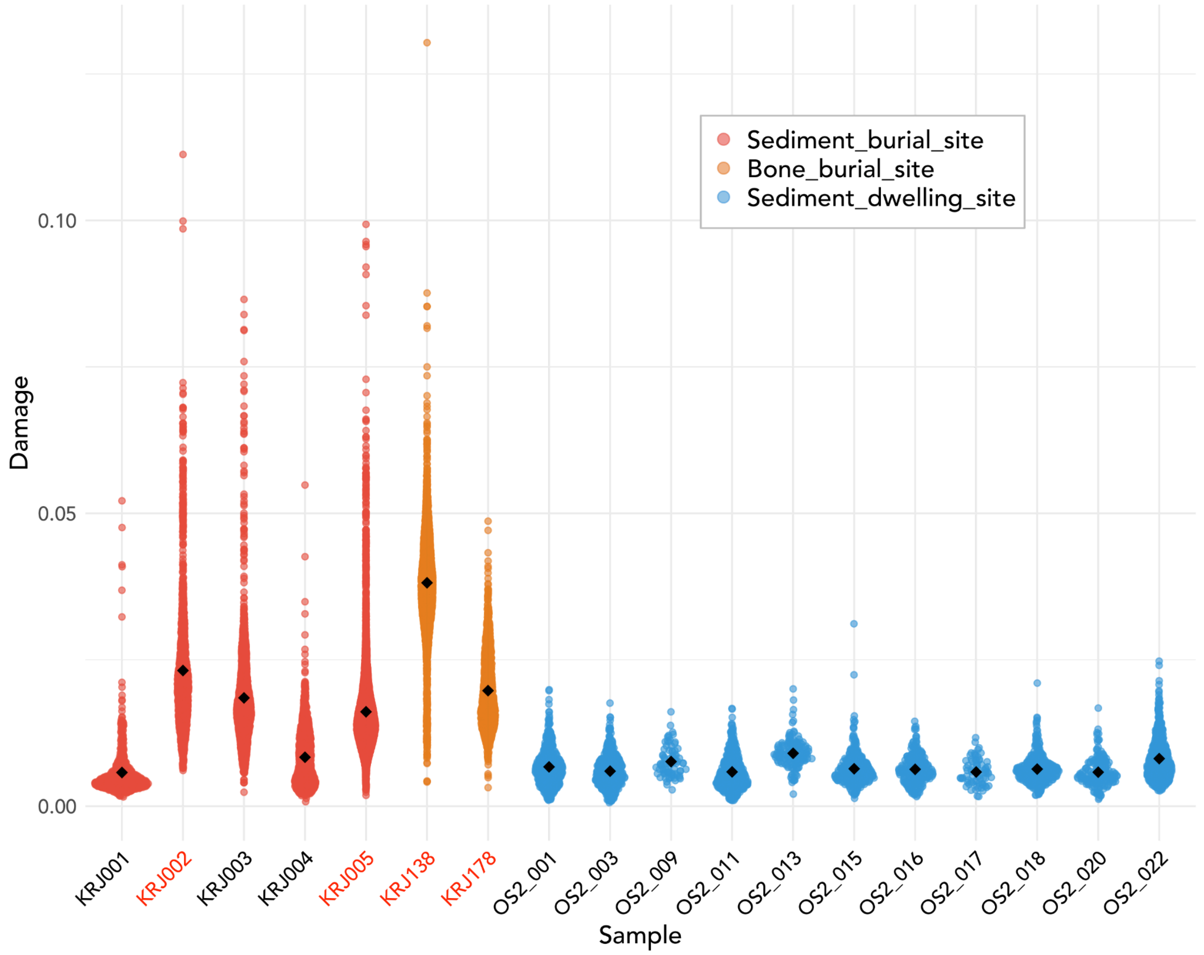


Figure S6. Damage degree of bacteria using shotgun datasets. Each dot represents the average DNA damage degree of each taxon. The black dot represents the average damage degree of all taxon. The red text color indicates samples that the haplogroup was determined from.
